# Concurrent *Ascaris* infection modulates host immunity resulting in impaired control of *Salmonella* infection in pigs

**DOI:** 10.1101/2024.02.21.581410

**Authors:** Ankur Midha, Larissa Oser, Josephine Schlosser-Brandenburg, Alexandra Laubschat, Robert M. Mugo, Zaneta D. Musimbi, Philipp Höfler, Arkadi Kundik, Rima Hayani, Joshua Adjah, Saskia Groenhagen, Malte Tieke, Luis E. Elizalde-Velázquez, Anja A. Kühl, Robert Klopfleisch, Karsten Tedin, Sebastian Rausch, Susanne Hartmann

**Author notes:** Address Correspondence to: Ankur Midha.

## Abstract

*Ascaris* is one of the most widespread helminth infections of humans and pigs, leading to chronic morbidity in humans and considerable economic losses in pig farming. Additionally, pigs are an important reservoir for the zoonotic bacterial pathogen *Salmonella,* where pigs can serve as asymptomatic carriers. Here, we investigated the impact of an ongoing *Ascaris* infection on the immune response to *Salmonella* in pigs. We observed higher bacterial burdens in experimentally coinfected pigs compared to pigs infected with *Salmonella* alone. *Ascaris-*infected pigs exhibited numerous hallmarks of a type 2 immune response in organs impacted by larval migration, including increased Th2 cells, increased IL-4 production, eosinophilia, and increased expression of CD206, a marker for alternatively activated macrophages. While we observed only mild changes in frequencies of CD4^+^ Treg, *Ascaris-* infected pigs had increased frequencies of CD8α^+^ Treg. We show that type 2 immune signals enhance susceptibility of macrophages to *Salmonella* infection *in vitro*. Furthermore, *Ascaris* impaired *Salmonella-*induced monocytosis and TNF-α production by myeloid cells. Hence, our data demonstrate widespread immunomodulation during an acute *Ascaris* infection that facilitates the microbial spread into gut-associated lymphoid tissue in a *Salmonella* coinfection.

**Importance:** In experimentally infected pigs we show that an ongoing infection with the parasitic worm *Ascaris suum* modulates host immunity to render pigs more susceptible to invading *Salmonella.* Both infections are widespread in pig production and the prevalence of *Salmonella* is high in endemic regions of human Ascariasis, indicating that this is a clinically meaningful coinfection. We observed a type 2 immune response to be induced during an *Ascaris* infection correlating with an increased susceptibility of pigs to the concurrent bacterial infection.

## Introduction

Helminth infections are widespread amongst humans and animals and occur alongside numerous microbial pathogens (1). *Ascaris lumbricoides* is the most prevalent helminth infection with over 800 million people infected globally (2). The closely related and zoonotic *Ascaris suum* is widespread in pig farming leading to significant production losses (3). Surveys of pig farms in different countries often find high infestation rates with most, if not all, farms impacted (4). *Ascaris* infection follows ingestion of eggs containing infective third-stage larvae (L3) which hatch in the host gut prior to migrating through host tissues by first invading the intestinal barrier, reaching the liver via the portal vein, and passing through the lungs 6-8 days post-infection (dpi) (5). The larvae then penetrate the alveoli, reaching the pharynx to be swallowed and return to the intestine where they reside and develop to L4 primarily in the jejunum from 8-10 dpi onwards (5).

Salmonellosis is one of the most commonly reported intestinal infections in the European Union, and is caused by serovars of *Salmonella enterica,* a zoonotic, facultative intracellular, food-borne bacterial pathogen (6). The prevalence of nontyphoidal *Salmonella* is high in helminth-endemic areas (7). Pigs are the second most common source of zoonotic *Salmonella* after poultry and *Salmonella* can be detected throughout the pig production chain (8, 9). Infection follows ingestion of contaminated food products, and once arriving in the small intestine, *Salmonella* can traverse the intestinal wall within minutes after invasion of microfold (M) and intestinal epithelial cells (10). Phagocytes, primarily macrophages, take up the invading bacteria and transport them to local gut-associated lymphoid tissues (11, 12). While symptomatic infections are usually accompanied by diarrhea and fever, pigs are often subclinical carriers with studies reporting up to 36% of pigs as asymptomatic shedders (13). In pigs, *Salmonella* can persist within macrophages in lymph nodes for months until market age (14) and the stress of transport can lead to bacterial egress and shedding, leading to increased *Salmonella* loads and contamination of work surfaces and food products (14, 15).

Helminth infections typically induce modified type 2 immune responses, characterized by the release of effector cytokines interleukin (IL-) 4, IL-5, and IL-13 from Th2 cells and type 2 innate lymphoid cells (ILC), and by heightened production of anti-inflammatory IL-10 by regulatory T (Treg) cells. This cytokine profile ensues in type 2 antibody responses, eosinophilia, the accumulation of alternatively-activated M2 macrophages, and in tissue remodelling including goblet cell hyperplasia designed to harm the migratory larval stages and to expel larval and adult worms from infected intestines (16). Accordingly, *Ascaris*-infected pigs have increased mRNA levels for Th2 and Treg-associated cytokines and there is evidence supporting a role for eosinophils in larval clearance (17–20). In contrast, immunity against *Salmonella* involves mixed type 1/3 responses carried by NK cells, Th1 cells, and Th17 cells and driven by pro-inflammatory cytokines such as tumor necrosis factor-α (TNF-α), IL-12, interferon-γ (IFN-γ), and IL-18, as well as IL-1β and IL17-A which support anti-bacterial responses by monocytes, conventionally-activated M1 macrophages, and neutrophils (21–24). Macrophages activated by pro-inflammatory cytokines can defend against intracellular bacteria; however, *Salmonella* can subvert the antimicrobial activity of macrophages and establish a niche within these cells by promoting alternative activation and preventing lysosome fusion while dwelling inside a *Salmonella*-containing vacuole (25–27).

Helminths have been shown to impair responses to bacterial antigens (28–31) and one study has shown increased *Salmonella* burdens during an ongoing helminth infection in mice (32). As such data are lacking in clinically-relevant, non-model animals, in this study we asked if an ongoing *Ascaris* infection impacts immune responses to *Salmonella* in pigs. We show that a concurrent *Ascaris* infection modulates host immunity to invasive *Salmonella* leading to increased bacterial burdens in coinfected pigs.

## Results

### *Ascaris*-Coinfected Pigs Have Elevated *Salmonella* Burdens

In an experimental coinfection we sought to determine if *Ascaris* infection impacts bacterial burdens in pigs. Commercial weaning pigs were infected with 4 consecutive inocula of 2000 *A. suum* eggs alone, 10^7^ CFU *S.* Typhimurium alone, both pathogens, or left uninfected (Figure 1A). Pigs in the coinfection group were infected first with *A. suum*, and then with *S.* Typhimurium 7 days later. In this experimental design, animals were dissected 14 days after the initial *A. suum* infection and 7 days after infection with *S.* Typhimurium. This time point was chosen as it represents the time when most *Ascaris* larvae have returned to the jejunum following tissue migration (5) and *Salmonella* have been carried into host tissues by phagocytes where they may persist long term (14, 33). Bacterial burdens were assessed locally in mLN of the jejunum and ileum (ileo-cecal mLN) and in the spleen to assess for systemic dissemination. At the primary site of *Salmonella* infection in the ileum, bacterial burdens were not significantly different, although two pigs in the coinfection group exhibited higher bacterial burdens than any of the pigs infected with *Salmonella* alone (Figure 1B). At the site of *Ascaris* infection in the jejunum, pigs infected with *Salmonella* alone were free of bacteria except for one pig while coinfected pigs had significantly elevated bacterial burdens (Figure 1B). None of the pigs had bacteria in the spleen suggesting that none developed a systemic infection (Figure 1B). None of the pigs in the trial developed any signs or symptoms of acute salmonellosis and weight gain was not impacted by either infection (Figure S1). Interestingly, coinfection did not appear to affect tissue pathology. Both *Ascaris-* and coinfected pigs demonstrated liver pathology due to migrating larvae whereas no significant pathology was seen in the lung (Figure S2). All infected pigs had similar pathology scores in the jejunum and ileum (Figure S2).

**Figure 1.**
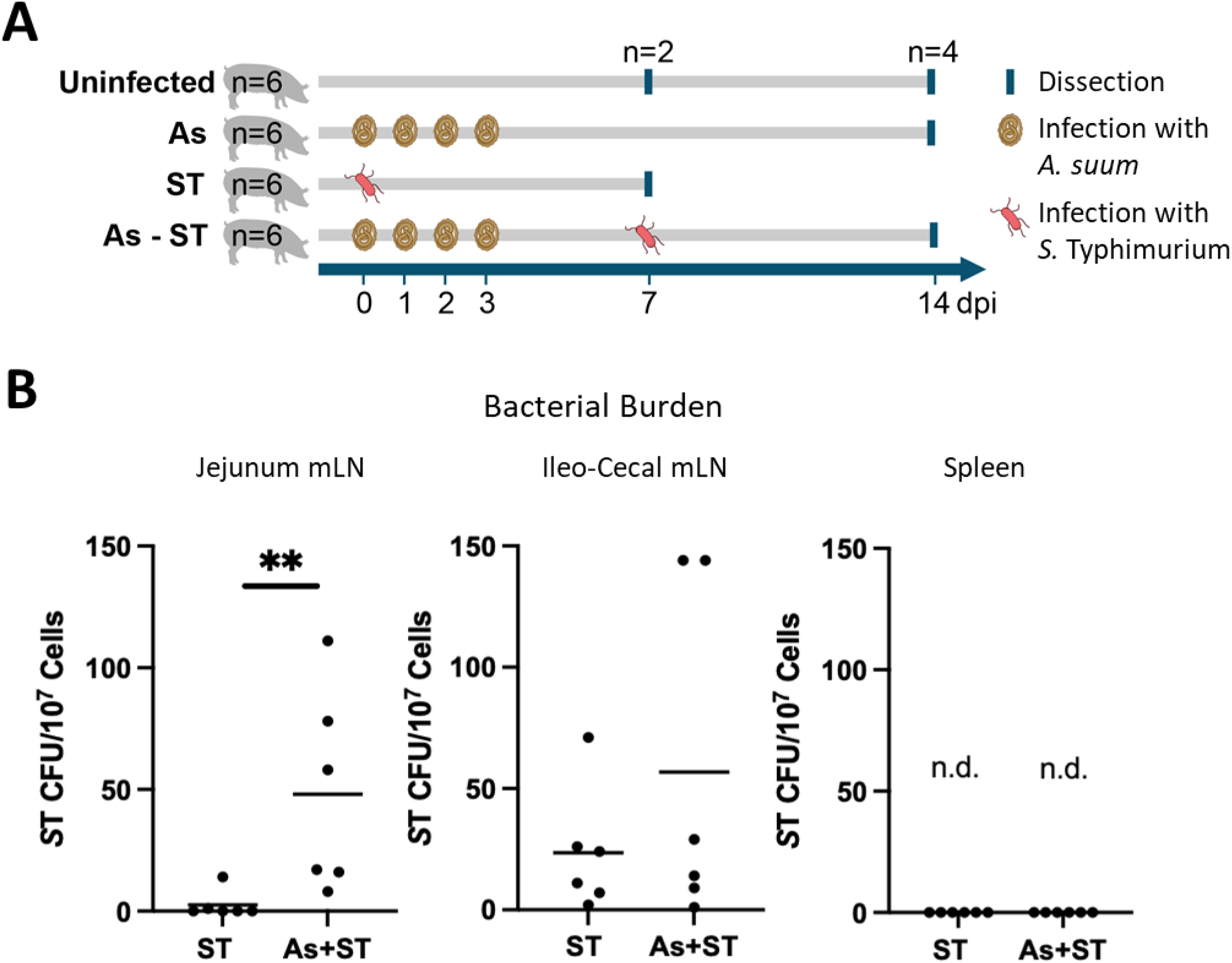
*Ascaris* and *Salmonella* are coinfecting pathogens of slaughterhouse pigs and experimental coinfection shows elevated *Salmonella* burdens. **A.** Experimental design. Crossbred six-week-old pigs (hybrid Landrace and Large White) were left uninfected (Ctrl), experimentally infected with 4 inocula of 2000 embryonated *A. suum* eggs (As), infected with 10^7^ colony forming units (CFU) *S.* Typhimurium (ST), or infected with both pathogens (As+ST). Pigs were dissected either 14 days post-infection (dpi) with *A. suum* or 7 dpi with *S.* Typhimurium and the same dissection timepoints were maintained in the coinfected group. **B.** *S.* Typhimurium CFU were assessed in the mesenteric lymph nodes (mLN) of the jejunum and ileum (ileo-cecal) and in the spleen; n.d. = not detected. Statistical significance determined by Mann-Whitney test, ** *p* ≤ 0.01.

### *Ascaris* Infection Induces a Modified Th2 Response and Suppresses Antibacterial T Cell Responses

Helminth infections typically induce a modified, IL-4-driven type 2 immune response characterised by Th2 and Treg cells which can impair immunity to bacterial and viral infections (1, 16). We therefore assessed type 2 T cell responses by quantifying the frequencies of GATA3^+^ (Th2 transcription factor) and IL-4^+^ (Th2 cytokine) T cells via flow cytometry (Figure 2A and 2B). In PBMC, only T cells from coinfected pigs exhibited significantly increased GATA3^+^ cells compared to uninfected controls (Figure 2A). However, a clear *Ascaris*-induced Th2 response could be seen as characterized by increased IL-4^+^ cells in PBMC of *Ascaris-* and coinfected pigs (Figure 2A). Interestingly, Th2 responses were not seen in the jejunum for any of the pigs (Figure 2B).

**Figure 2.**
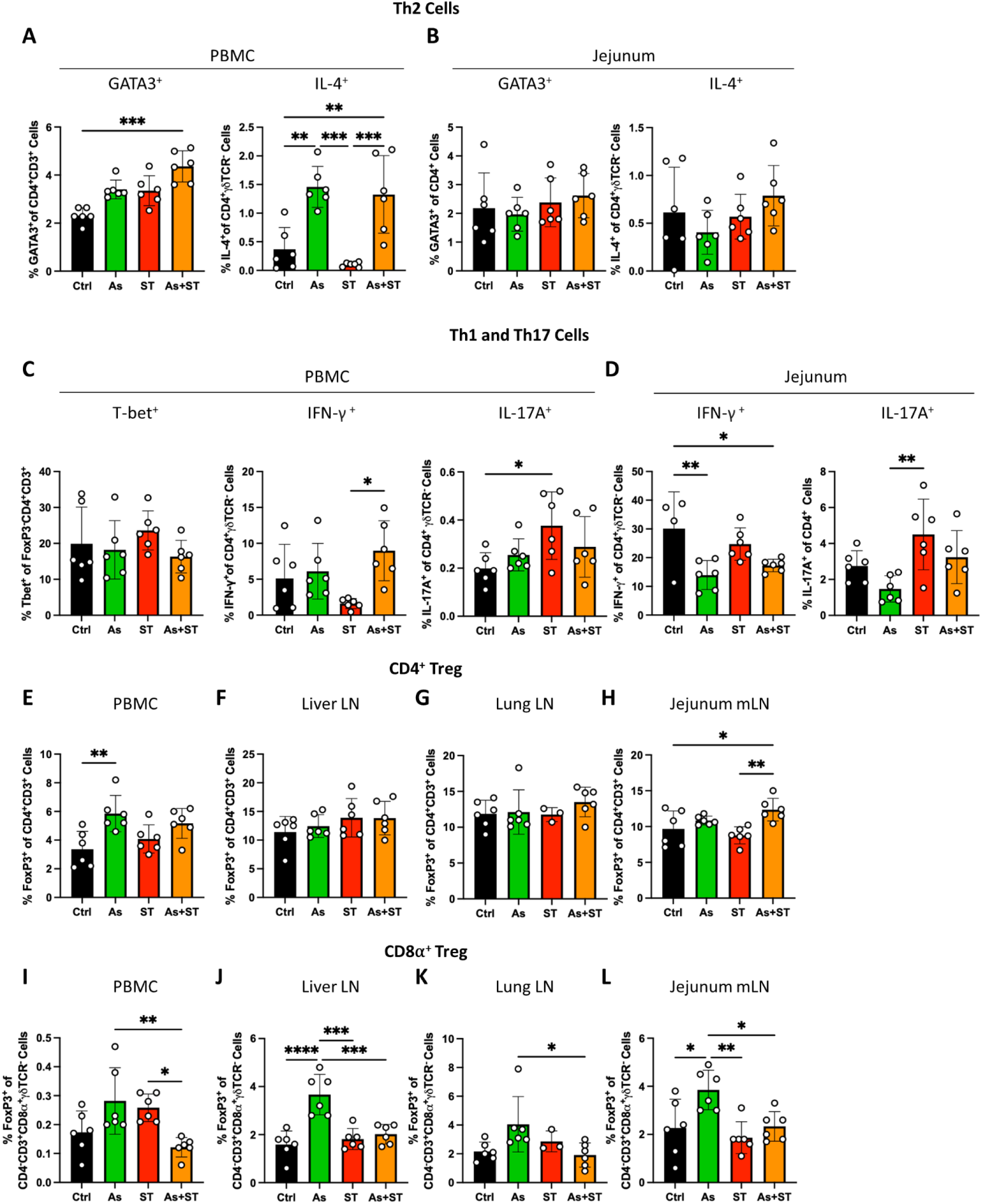
*Ascaris* infection induces a modified type 2 T cell response. **A.** Th2 cell subset frequencies in PBMC. GATA3 transcription factor expression was assessed in live CD4^+^CD3^+^ cells while IL-4 cytokine expression was assessed in live CD4^+^CD3^+^γδTCR^-^ cells. **B.** Th2 cell subset frequencies in jejunum. **C.** Th1 and Th17 cell subset frequencies in PBMC. T-bet transcription factor expression was assessed in live CD4^+^CD3^+^ cells while IFN-γ, and IL-17A cytokine expression was assessed in live CD4^+^CD3^+^γδTCR^-^ cells. **D.** Th1 and Th17 cell subset frequencies in jejunum. **E.** CD4^+^ Treg frequencies (FoxP3^+^CD4^+^CD3^+^) in PBMC. **F.** CD4^+^ Treg frequencies in liver LN. **G.** CD4^+^ Treg frequencies in lung LN. **H.** CD4^+^ Treg frequencies in jejunum mLN. **I.** CD8α^+^ Treg frequencies (FoxP3^+^CD4^-^CD3^+^CD8α^+^γδTCR^-^) in PBMC. **J.** CD8α^+^ Treg frequencies in liver LN. **K.** CD8α^+^ Treg frequencies in lung LN. **L.** CD8α^+^ Treg frequencies in jejunum mLN. Statistical significance was determined by one-way analysis of variance followed by Tukey’s multiple comparisons test, * *p* ≤ 0.05, ** *p* ≤ 0.01, *** *p* ≤ 0.01, **** *p* ≤ 0.001.

In contrast to helminths, type 1 and type 3 immune responses characterised by the differentiation and expansion of IFN-γ^+^ Th1 and IL-17a^+^ Th17 cells are protective against intracellular pathogens including *Salmonella* (9, 21). We therefore assessed inflammatory T cell responses by quantifying the frequencies of T-bet^+^ (Th1 transcription factor), IFN-γ^+^, and IL-17A^+^ T cells (Figure 2C and 2D). As shown in Figure 2C, while T-bet expression was similar in CD4^+^ PBMC of all groups, cells from *Salmonella* infected pigs homogenously displayed the lowest IFN-γ production, paired with relatively high IL-17A responses by PBMC (Figure 2C). Th1 and Th17 responses did not differ between either *Ascaris* single or coinfected pigs and the naïve controls (Figure 2C). In the jejunum, *Ascaris-* and coinfected pigs exhibited lower levels of IFN-γ^+^ T cells compared to uninfected controls, whereas *Salmonella-*infected, but not coinfected pigs, had higher levels of IL-17A^+^ T cells (Figure 2D).

In order to limit tissue damage, regulatory cells including Treg can suppress inflammatory responses (34). While advantageous in limiting immunopathology, Treg induced or expanded during infection can limit inflammatory responses necessary for pathogen clearance (34). Helminths, including *Ascaris*, have been shown to induce CD4^+^ Treg which can restrain anti-parasitic type 2 responses (35, 36). Therefore, we assessed for regulatory responses by quantifying FoxP3^+^ (Treg transcription factor) T cells in systemic circulation and in the lymph nodes draining the organs impacted by *Ascaris* tissue migration (Figure 2E-H). *Ascaris-*infected pigs exhibited increased CD4^+^ Treg in PBMC (Figure 2E) and in the jejunum mLN (Figure 2H). Interestingly, no CD4^+^ Treg inductions were seen in the lymph nodes of the liver and lung (Figure 2F and 2G).

While less frequently studied than CD4^+^ Treg, FoxP3^+^ as well as FoxP3^-^ CD8^+^ Treg can also be induced by various infections (34, 37). The murine helminth *Heligmosomoides polygyrus* induces robust CD8^+^ Treg capable of controlling colitis (38) and preventing autoimmune diabetes (39). We therefore quantified frequencies of CD8α^+^ Treg (Figure 2I-L). Interestingly, while there were very low levels found in systemic circulation, we found elevated CD8α^+^ Treg in the lymph nodes of organs impacted by *Ascaris* larval tissue migration (liver, lung, jejunum) in *Ascaris*-infected pigs (Figure 2J-L). In contrast to CD4^+^ Treg, there was no indication of CD8α^+^ Treg induction in coinfected pigs.

Taken together, *Ascaris* infection induced a modified type 2 response characterised by Th2 cells and dominant CD4 as well as CD8α Treg cells while pigs responded to *Salmonella* infection with relatively prominent systemic and local Th17 cell responses. *Ascaris* infection created a modulatory immune environment wherein antibacterial IL-17^+^ T cell responses against *Salmonella* showed a trend of being attenuated.

### *Ascaris* infection modulates the host granulocyte response

As adaptive immune responses are a product of early innate immune signals, we turned our attention to innate immune responses. *Ascaris* and *Salmonella* differentially induce granulocytes, common first-responders in different infections. Tissue-migrating *Ascaris* larvae are associated with increased levels of eosinophils which may be protective (19, 20) whereas neutrophils play a considerable role in clearing *Salmonella* (22). Therefore, we sought to quantify granulocytes in tissues impacted by the two pathogens. First, we quantified neutrophils in blood smears and broncheoalveolar lavage (BAL) cytospin preparations (Figure 3A). Interestingly, we did not see a systemic induction of neutrophils in *Salmonella-* or coinfected pigs (Figure 3A). However, there appeared to be a decrease in mean neutrophil frequencies in the blood of *Ascaris*-infected pigs (22.8 ± 3.7% vs. 34.5±7.5% in uninfected controls), though this difference was not statistically significant (Figure 3A). Curiously, coinfected pigs displayed higher levels of neutrophils in the BAL, while either pathogen alone had no influence on neutrophils in this compartment (Figure 3A). We also assessed the ability of *Ascaris* to modulate neutrophil recruitment by assessing *IL8* mRNA levels and found that *Ascaris-* and *Salmonella-*infected pigs had decreased *IL8* mRNA expression in the lung compared to uninfected controls and coinfected pigs (Figure 3B). We then quantified eosinophil frequencies which were unremarkable in the blood (Figure 3C). In contrast, eosinophil frequencies in the BAL were markedly increased in both *Ascaris-* and coinfected pigs (Figure 3C). To further assess the impact of *Ascaris* on eosinophils, we quantified eosinophils by histology in the intestine and found that eosinophils were markedly increased in the jejunum and ileum of *Ascaris-*and coinfected pigs (Figure 3D). Thus, consistent with previous findings, *Ascaris* infection resulted in eosinophilia; hence, *Ascaris-* and coinfected pigs demonstrated a type 2 bias in the innate granulocyte response which may compromise innate responses against incoming *Salmonella*.

**Figure 3.**
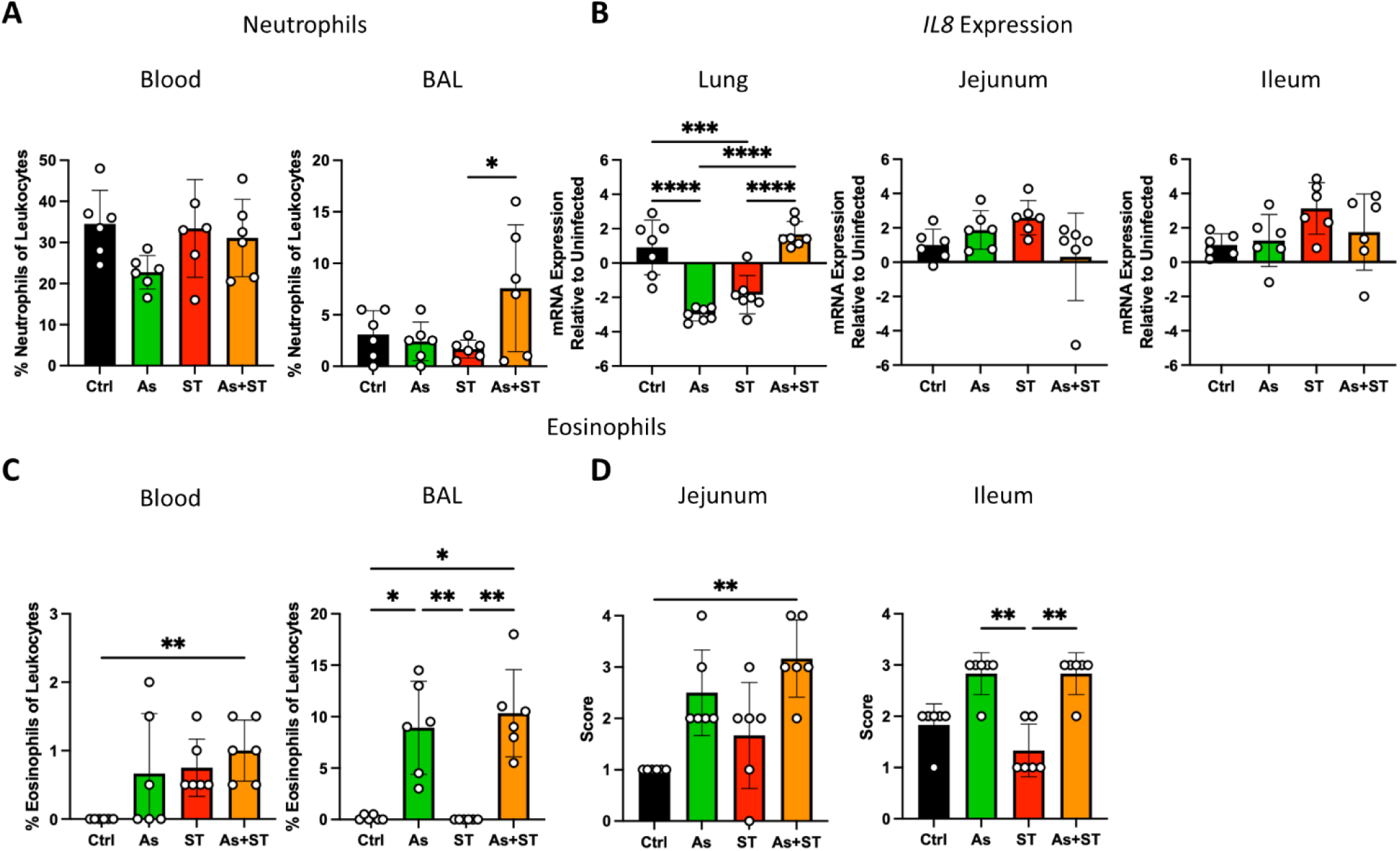
*Ascaris* infection modulates granulocyte responses. **A.** Neutrophil counts in blood smears (left) and broncheoalveolar lavage (BAL) cytospins (right) as a percentage of leukocytes. **B.** Bar graphs representing mean relative mRNA expression (±SD) of *IL8* in lung, jejunum, and ileum. **C.** Eosinophil counts in blood smears (left) and BAL cytospins (right) as a percentage of leukocytes. **D.** Tissue eosinophil counts by histological scoring of different tissue sites as described in the methods. Data were tested for normality and statistical significance was determined by one-way analysis of variance followed by Tukey’s multiple comparisons test (Neutrophils in blood & BAL, *IL8* mRNA expression) or Kruskal-Wallis test followed by Dunn’s multiple comparisons test (all other plots), * *p* ≤ 0.05, ** *p* ≤ 0.01, *** *p* ≤ 0.001, **** *p* < 0.0001.

### *Ascaris* Infection Decreases Macrophage Frequencies and Increases M2 Macrophage Marker Expression

As macrophages are a favoured site for *Salmonella* persistence (24, 26), we next assessed the impact of *Ascaris* infection on this cell type. We first assessed the frequencies of macrophages by quantifying CD172a^+^CD163^+^CD203a^+^ cells (40) in relevant tissues (Figure 4A). Macrophage frequencies remained unchanged between groups in the liver (Figure 4B). Interestingly, *Ascaris* infection significantly reduced tissue macrophage frequencies in the lung, jejunum, and ileum (Figure 4B).

**Figure 4.**
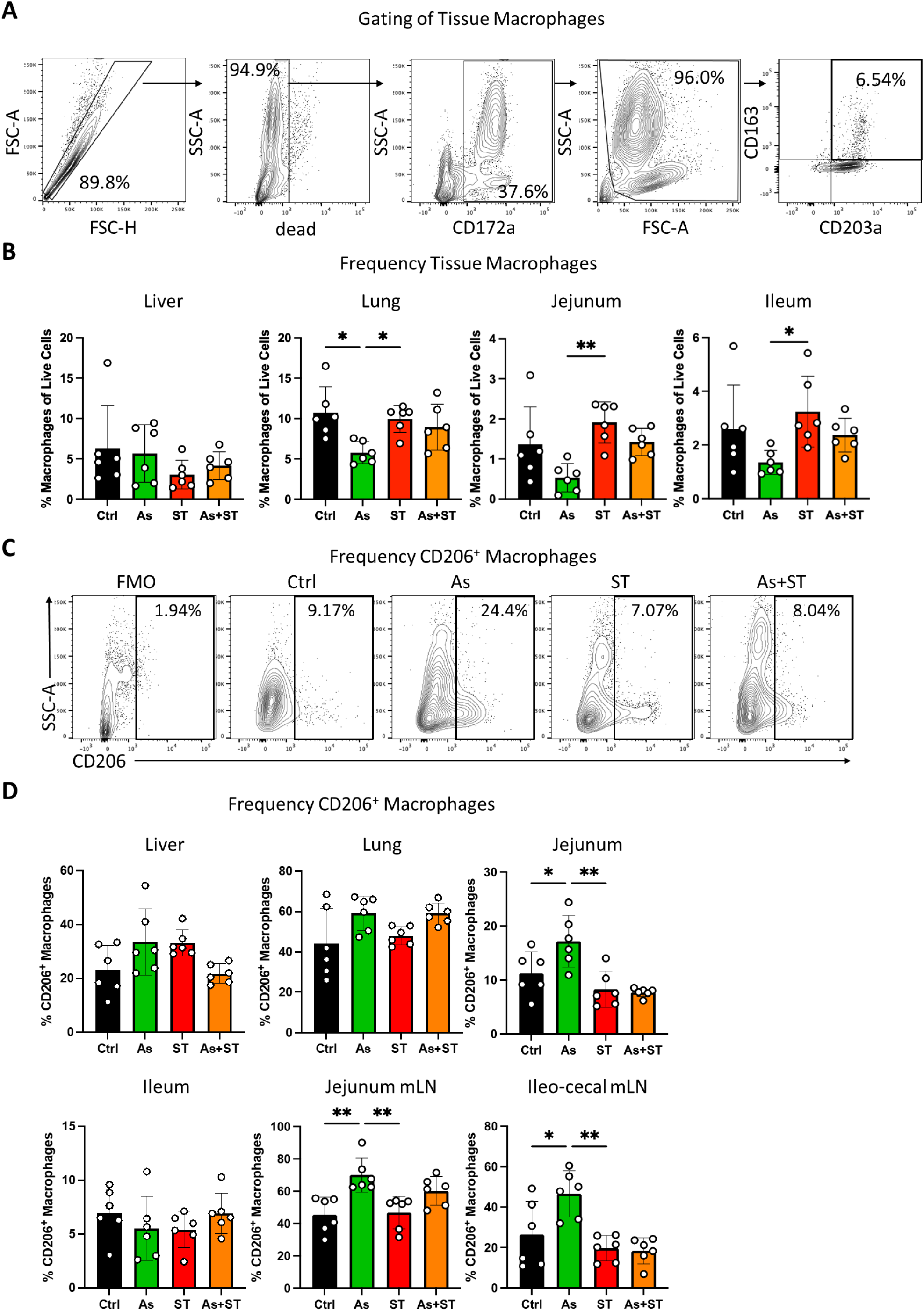
*Ascaris* infection reduces macrophage frequencies and induces M2 polarization. **A.** Gating strategy to identify tissue macrophages, defined as live CD172a^+^CD163^+^CD203a^+^ cells. Plots report the gating of jejunum lamina propria macrophages. **B.** Macrophage frequencies in liver, lung, jejunum lamina propria, and ileum lamina propria. **C.** Representative flow cytometry plots of CD206^+^ macrophages in jejunum lamina propria. **D.** Frequencies of CD206^+^ macrophages in different tissues. Data were tested for normality and statistical significance was determined by one-way analysis of variance followed by Tukey’s multiple comparisons test, * *p* ≤ 0.05, ** *p* ≤ 0.01.

Previous work has shown that macrophage polarization state impacts *Salmonella* growth and persistence; M1 macrophages suppress bacterial growth while M2 macrophages are associated with proliferating bacteria (27). The mammalian mannose receptor (CD206) is expressed by several types of tissue macrophages and is considered an M2 macrophage marker as its expression is promoted by IL-4 and increased during helminth infections (40, 41). We therefore hypothesized that *Ascaris* infection would increase M2 macrophage polarization. To test this, we assessed frequencies of CD206^+^ macrophages in tissues where macrophage frequencies had been impacted by *Ascaris* infection (Figures 4C and 4D). Frequencies of CD206^+^ macrophages were unchanged in the liver and ileum (Figure 4D). In the lung, while not statistically significant, the frequency of CD206^+^ cells tended to increase in all infected pigs relative to uninfected controls (Figure 4D). In the jejunum, *Ascaris* infection increased the frequency of CD206^+^ cells (Figures 4C and 4D). As *Salmonella* can persist within macrophages in gut-associated lymphoid tissue (11, 12), we also assessed the frequency of CD206^+^ macrophages in intestinal mLN. Remarkably, *Ascaris-*infected pigs had more CD206^+^ macrophages in both mLN sites indicating that macrophages at the site of *Salmonella* persistence are modulated by *Ascaris* infection. Together, these data suggest that *Ascaris* infection reduces tissue macrophage populations while increasing M2 polarization in remaining macrophages, thereby increasing susceptibility to invading *Salmonella*.

### *Ascaris*-Induced Immune Environment Enhances Macrophage Susceptibility to *Salmonella*

We next aimed to determine whether the *Ascaris-*induced immune environment had an impact on intracellular growth of *Salmonella.* We tested this by exposing alveolar macrophages from uninfected pigs to different cytokine treatments (Figure 5A). Macrophages were either left untreated or treated with M1-polarizing IFN-γ, or M2-polarizing IL-4 and IL-13. Then, macrophages were infected with GFP-expressing *Salmonella* and assessed for bacterial growth by fluorescence microscopy and flow cytometry. Macrophages treated with IFN-γ showed a reduction in intracellular *Salmonella* while M2 signals increased bacterial burdens (Figure 5B-D). Thus, pro-inflammatory signaling (IFN-γ) is protective against *Salmonella* while helminth-induced signaling (IL-4/13) impairs macrophage antibacterial activity, leading to higher intracellular bacterial burdens.

**Figure 5.**
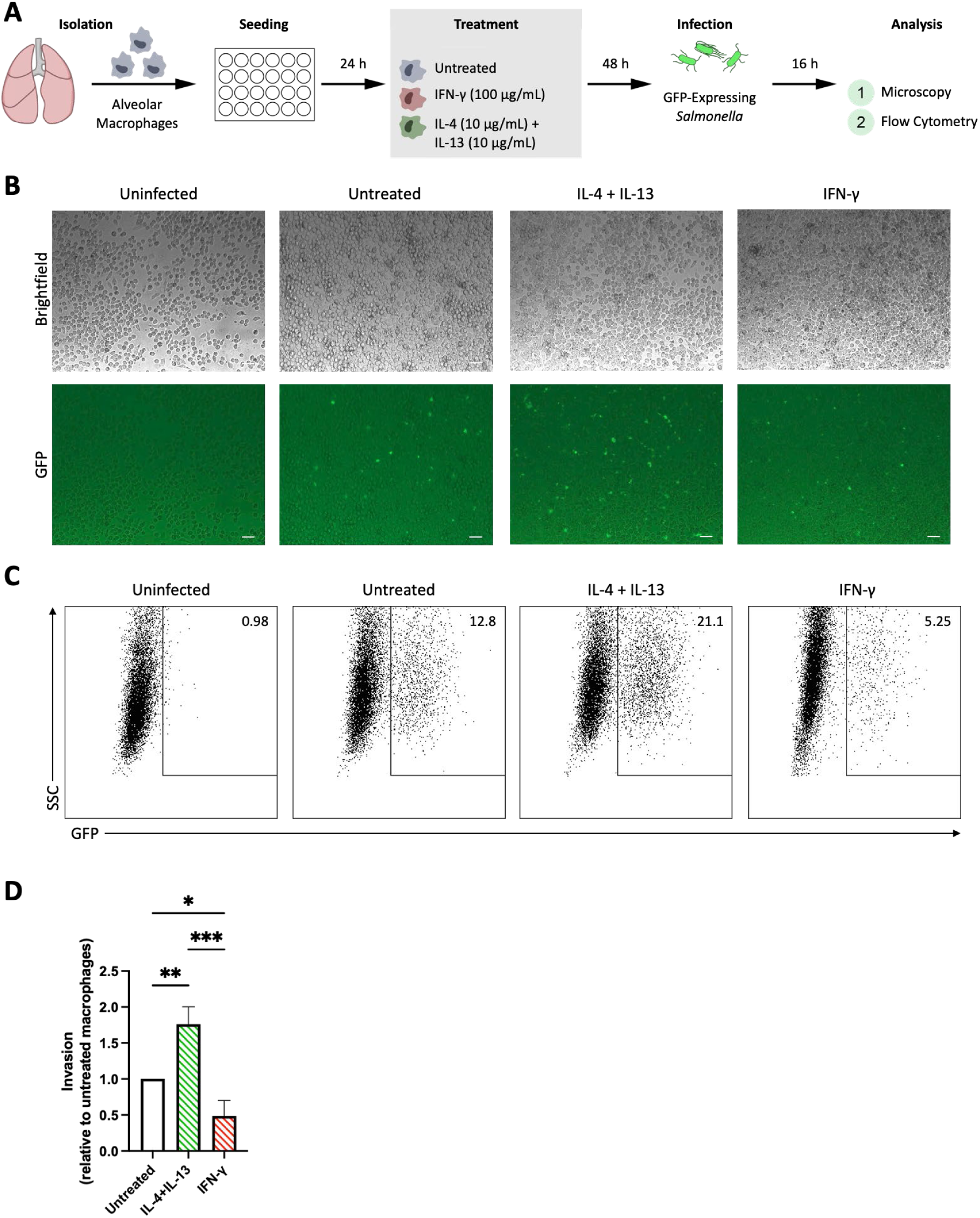
*Ascaris*-modulated immune environment enhances macrophage susceptibility so *Salmonella* infection. **A.** Experimental overview. Alveolar macrophages were isolated from the lungs of uninfected pigs. Macrophages were cultured in culture media with and without cytokine treatments for 48 h prior to infection. Macrophages were infected at a multiplicity of infection of 10 with GFP-expressing *Salmonella* Typhimurium. Sixteen hours later, cells were assessed for bacterial burden by fluorescence microscopy and flow cytometry. **B.** Representative images of GFP-expressing *Salmonella* infection of macrophages. Cells visualized with 20x objective. Data are representative of three independent experiments. **C.** Representative flow cytometry plots of GFP^+^ macrophages. Data are representative of three independent experiments. **D.** Columns represent mean invasion (with untreated macrophages set to 1.0) from three independent experiments + standard deviation. Statistical significance was determined by one-way analysis of variance followed by Tukey’s multiple comparisons test, * *p* ≤ 0.05, ** *p* ≤ 0.01, *** *p* ≤ 0.01.

### *Ascaris* Infection Impairs Monocyte Responses and Reduces Monocyte Frequencies

In addition to evidence for alternative activation of macrophages by *Ascaris* infection, we also observed reduced macrophage frequencies in the mucosal tissues of lung, jejunum, and ileum, possibly indicating the impaired recruitment of monocytes from the blood; monocytes can be recruited to peripheral sites and differentiate into macrophages in peripheral tissues during infection (42). Furthermore, previous studies have shown that monocytes participate in *Salmonella* clearance (9, 21–23). We therefore assessed the impact of both pathogens on monocyte frequencies by flow cytometry. *Salmonella* infection was associated with increased monocyte frequencies in the blood (Figure 6A). In contrast, coinfected pigs did not exhibit elevated monocytes and mean monocyte frequencies were lower in PBMC of *Ascaris-* (2.4 ± 0.6%) and coinfected (1.6 ± 0.7%) pigs compared to uninfected controls (3.2 ± 1.9%) (Figure 6A). Next, we assessed the responsiveness of monocytes to *ex vivo* stimulation with IL-12 and IL-18. Remarkably, while monocytes from *Salmonella*-infected pigs produced TNF-α in response to cytokine stimulation, monocytes from *Ascaris-* and coinfected pigs did not (Figure 6B and 6C). We then sought to determine if in addition to reduced tissue macrophages and blood monocytes, the overall population of monocytes and macrophages was reduced at the site of infection in the intestine. We did so by assessing FSC^high^CD172a^+^ cells in the gut (Figure 6E). Remarkably *Ascaris* infection reduced the overall pool of monocytes and macrophages in both the jejunum and the ileum, while *Salmonella* increased monocyte-macrophage frequencies in the ileum (Figure 6F). Interestingly, coinfected pigs did not display an increase in ileal monocyte-macrophage frequencies (Figure 6F). Finally, we asked if infection altered intestinal expression of monocyte chemoattractant protein-1 (MCP-1), also known as CCL2. We assessed mRNA levels of *CCL2* and did not observe a demonstrable impact of either infection on *CCL2* levels in the jejunum, while coinfected pigs had increased *CCL2* mRNA levels in the ileum (Figure 6G). Together, these data demonstrate a considerable blunting of monocyte responses against *Salmonella* by *Ascaris* and an inability of the host to recruit monocytes to the site of infection to replenish monocyte and macrophage populations decreased during *Ascaris* infection. Thus, *Ascaris*-infected pigs are compromised in their response to invading *Salmonella*.

**Figure 6.**
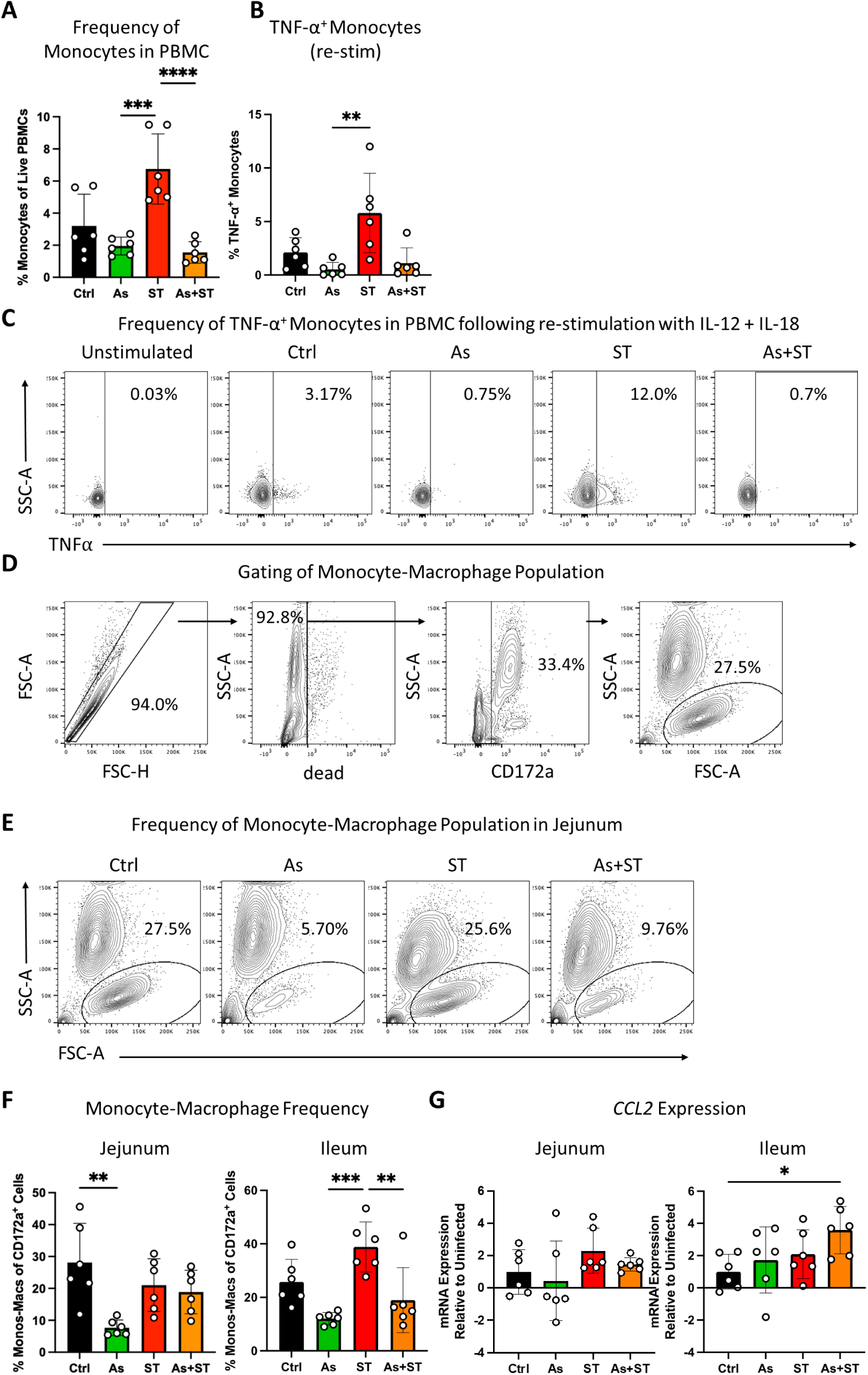
*Ascaris* infection blunts *Salmonella*-induced monocytosis and inflammatory cytokine secretion. **A.** Monocyte frequencies in porcine PBMC. Monocytes were defined as live FSC^high^CD3^-^CD8^-^CD16^+^ cells. **B.** Monocyte responses assessed by TNF-α expression in monocytes stimulated with IL-12 + IL-18. **C.** Representative flow cytometry plots of re-stimulated monocytes in PBMC. **D.** Gating strategy to identify tissue monocytes-macrophages, defined as live CD172a^+^FSC^high^ cells. Shown here: jejunum lamina propria monocytes-macrophages. **E.** Representative flow cytometry plots of monocyte-macrophage population in the jejunum lamina propria. **F.** Frequencies of monocytes-macrophages in the intestine. **G.** Bar graphs representing mean relative mRNA expression (±SD) of *CCL2* in jejunum and ileum. Data were tested for normality and statistical significance was determined by one-way analysis of variance followed by Tukey’s multiple comparisons test (Monocyte frequencies, monocyte-macrophage frequencies, mRNA expression) or Kruskal-Wallis test followed by Dunn’s multiple comparisons test (TNF-α expression), * *p* ≤ 0.05, ** *p* ≤ 0.01, *** *p* ≤ 0.01, **** *p* ≤ 0.001.

**Figure 7.**
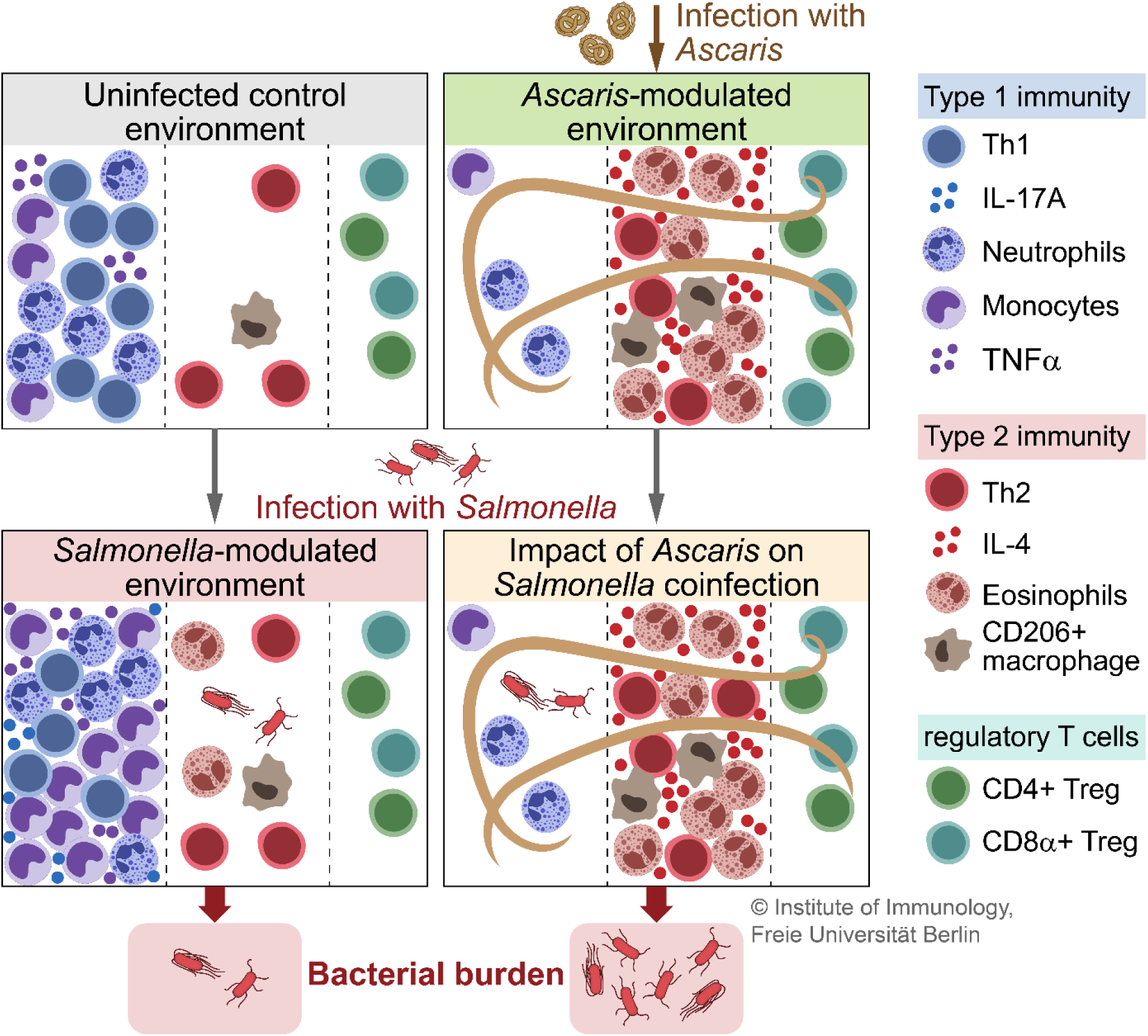
*Ascaris* infection modulates host immunity creating an immune environment more permissive for invading *Salmonella* (graphical abstract). *Ascaris* and *Salmonella* induce opposing immune responses. During an ongoing *Ascaris* infection, invading *Salmonella* encounter fewer neutrophils, monocytes, and macrophages. The remaining monocytes and macrophages are less responsive to *Salmonella* and exist within a type 2 immune environment (more IL-4, less pro-inflammatory cytokines). Collectively, an *Ascaris-*modulated immune environment allows *Salmonella* to establish itself, leading to higher bacterial burdens in coinfected pigs.

## Discussion

Helminths are amongst the most prevalent infectious agents globally and there are ample evidence indicating increased susceptibility to co-endemic microbial pathogens in helminth-infected hosts (1, 32). Here, we show that acute *Ascaris* infection modulates host immunity rendering the host more susceptible to infection with *Salmonella*. To the best of our knowledge, this is the first study to assess and compare porcine immune responses across multiple cell types against *Salmonella* and *Ascaris* coinfections, and to demonstrate a clinically meaningful interaction between the two pathogens. Our data are indicative of an *Ascaris*-modulated immune environment that is less able to deal with incoming *Salmonella*.

*Ascaris-* and coinfected pigs demonstrated a helminth-modulated T cell response characterised by increased Th2 cells and Treg, as well as reduced IFN-γ and IL-17A producing T cells (Figure 2). Interestingly, we observed differential Treg responses, wherein classical CD4^+^ Tregs were either not induced during infection or induced in both *Ascaris-* and coinfected pigs, contrasting with a rise in CD8^+^FoxP3^+^ Treg seen only in *Ascaris-*infected pigs. While CD4^+^ Tregs are known to modulate type 2 immunity (35, 36), CD8^+^ Treg are less well studied in helminth infections. Shimokawa and colleagues demonstrated that the disaccharide trehalose, produced by the murine helminth *H. polygyrus,* supported the growth of intestinal *Ruminococcus* spp. which in turn induced potently immunosuppressive CD8^+^ Treg in mice (39). *Ascaris* can also produce trehalose (43) and we have previously reported *Ruminococcus* to be a member of *Ascaris’* own intestinal microbiome (44). Future studies should assess the impact of acute *Ascaris* and *Salmonella* infections on intestinal microbial communities and metabolites to determine their influence on observed immune responses.

Neutrophils are early responders during *Salmonella* infection and mediate bacterial clearance (22, 45). In our study, we did not see neutrophilia in *Salmonella-*infected pigs; however, Burdick Sanchez and colleagues have shown that neutrophil levels peak in the first 24 h post-infection with *S.* Typhimurium before declining (45). Thus, by 7 dpi in our study we would not expect to see increased levels of neutrophils. Interestingly, we found that *Ascaris* infection reduced neutrophil levels (Figure 3A) indicating that incoming *Salmonella* have fewer neutrophils to contend with in *Ascaris-*infected hosts. Consistent with prior studies (19, 20), *Ascaris*-and coinfected pigs exhibited eosinophilia. During helminth infections, eosinophils support type 2 immunity by secreting IL-4, IL-5, IL-10, and IL-13 (reviewed in (46)). However, eosinophils can also secrete classical Th1 cytokines including IFN-γ and TNF-α (47). Nevertheless, while human eosinophils are indeed phagocytic, they are apparently less bactericidal than neutrophils (48). Our data indicate that the impact of *Ascaris* on host granulocytes renders hosts more susceptible to incoming *Salmonella,* although neutrophils eventually recover in coinfected hosts. Further studies are required to assess cytokine production and bactericidal activity of granulocytes in *Ascaris*-infected hosts.

Chen and colleagues reported monocyte recruitment to the lung and differentiation into M2-like alveolar macrophages during *Nippostrongylus brasiliensis* infection in mice (49). In contrast, our monocyte and macrophage data do not show similar tendencies in pigs based on monocyte and tissue macrophage frequencies. In agreement with Masure et al. who observed a decrease in macrophage counts in the cecum of *Ascaris*-infected pigs (20), we observed decreased macrophage frequencies in the jejunum and ileum during acute ascariasis (Figure 4A). One study reported that ES from the liver fluke *Fasciola hepatica* induced apoptosis of murine peritoneal macrophages (50). Thus, further studies are warranted to determine if *Ascaris* infection induces macrophage death. While we found evidence for M2 macrophage polarization in *Ascaris*-infected pigs, macrophages in coinfected pigs are similar to those in *Salmonella*-infected pigs (Figure 4D). However, macrophages have been shown to be highly plastic (51). Nevertheless, our data from *in vitro*-treated alveolar macrophages indicates that *Ascaris*-induced immune signals compromise the ability of macrophages to suppress intracellular *Salmonella* growth (Figure 5).

In a coinfection study in mice with *H. polygyrus* and *S.* Typhimurium, Rückerl et al. found that blood monocyte-derived macrophages could displace tissue resident macrophage population (51). We observed a drastic impairment of host monocyte responses in *Ascaris*- and coinfected hosts (Figure 6). In addition to neutrophils, monocytes are also recruited during salmonellosis and porcine monocytes can rapidly suppress intracellular *Salmonella* (23, 52). While monocytosis was observed in response to *Salmonella*, coinfected pigs did not exhibit elevated monocyte levels (Figure 6A). Furthermore, monocytes from *Ascaris-* and coinfected pigs did not produce TNF-α in response to cytokine stimulation, unlike their counterparts from *Salmonella*-infected pigs (Figure 6B). This is consistent with previous work from our laboratory demonstrating ablated LPS-mediated TNF-α production by monocytes following treatment with *A. suum* excretory/secretory products (ES) (53). Similarly, Almeida et al. observed *A. suum* ES-mediated suppression of TNF-α responses in LPS-treated human monocyte-derived macrophages (54). Thus, acutely *Ascaris-*infected pigs may have fewer monocytes and macrophages present to fight invading pathogens, and those that are present are compromised in their ability to suppress bacterial growth.

While we observed increased *Salmonella* burdens in pigs with a concurrent *Ascaris* infection consistent with data from mice infected with *H. polygyrus* and *S.* Typhimurium, Reynolds and colleagues found that *Salmonella* coinfection was promoted by a helminth-modulated intestinal metabolome (32). Intestinal microbes not only produce bioactive metabolites, they also contribute to colonisation resistance to protect from various infections, including salmonellosis (55). We and others have shown that *Ascaris* infection alters the host gut microbiome (44, 56–58). Furthermore, *Ascaris* has been reported to produce acetate (59) which has been shown to enhance invasion of avian intestinal epithelial cells by *S.* Enteritidis (60). The contribution of an *Ascaris-*modulated microbiome and metabolome to susceptibility to microbial infection is an exciting area for further research.

Limitations of this study arose from the study design. Assessing bacterial burdens at 7 dpi only allows an assessment of the pathogenesis of infection between acute salmonellosis which peaks around 2-3 dpi (61) and persistence. Extended studies are required to assess bacterial burdens weeks and months after infection. Similarly, we have likely missed the expected neutrophilia during salmonellosis due to the experimental timeline. An intriguing question not answered by our study design is whether *Salmonella* can modulate host responses against *Ascaris.* A previous study by Steenhard et al. in which pigs were first infected with *S.* Typhimurium, then infected with 2500 *A. suum* eggs twice weekly for seven weeks found no impact on *Salmonella* burdens when *Ascaris* was introduced starting at 3 dpi and the authors reported no impact of *Salmonella* on worm burdens (62). Importantly, immune parameters were not studied in detail in this study. Neutrophils and macrophages have been shown to be modulated by *Salmonella*, allowing intracellular survival (27, 63). *Salmonella*-infected pigs also exhibit a disrupted intestinal microbiome (64, 65). Thus, whether *Salmonella* might influence ascariasis directly through its interaction with immune cells or indirectly via the microbiome is worthy of further study.

Collectively, our observations demonstrate that *Ascaris* is capable of immunomodulation even during acute infection. This modulation establishes an immune environment permissive for infection with other pathogens (summarized in Figure 9). Our results in a clinically relevant model suggest that helminth infections should be more closely monitored during pig production. Further studies on other coinfecting pathogens of interest, such as *Campylobacter jejuni, Streptococcus suis,* influenza, and porcine reproductive and respiratory syndrome viruses is warranted.

## Materials and Methods

### Experimental infection of pigs

The animal experiment was performed at Freie Universität Berlin under the principles outlined in the European Convention for the Protection of Vertebrate Animals used for Experimental and other Scientific Purposes and in accordance with the German Animal Welfare Act. Ethical approval was granted by the Berlin State Office of Health and Social Affairs (Landesamt für Gesundheit und Soziales; approval number G0212/20).

A total of 24 weaning hybrid (German landrace and large white) pigs of both sexes were obtained from a conventional breeder (Brandenburg, Germany) at 6 weeks of age. The pigs were allowed to acclimate for 11 days prior to the experiment. Water was given ad libitum and food was provided twice daily according to weekly body weight measurements. Wood chips were used as bedding and straw and toys were provided for enrichment. The pigs were randomly assigned to four groups, balanced for body weight and sex. The four groups included uninfected controls (Ctrl, n=6), pigs infected with *A. suum* (As, n=6), pigs infected with *S.* Typhimurium (ST, n=6), and animals infected with both pathogens (As+ST, n=6).

Infective *A. suum* eggs were collected and prepared as previously described (44). Adult female worms obtained from a local slaughterhouse were cultured overnight at 37 °C. Eggs released into the culture medium were collected, washed, and incubated at room temperature in the dark for 8 weeks. Pigs were orally infected with 2000 embryonated *A. suum* eggs/day for 4 days. The inoculum was fed to the pigs on store-bought waffles.

*S.* Typhimurium definitive type 104 (strain BB440) is a nalidixic acid resistant zoonotic pathogen and was used because it was originally obtained from a pig with acute salmonellosis (14). *Salmonella* were grown in Luria-Bertani (LB) broth (Carl Roth GmbH, Karlsruhe, Germany) with aeration at 37 °C to an optical density at 600 nm of 1.5-2.0. Pigs were orally inoculated with 10^7^ colony forming units (CFU) delivered on waffles, as for the *A. suum* inoculum. Pigs were closely monitored (body weight, visual observation of pigs and their feces) in the days following infection.

Dissection timepoints were 14 dpi for *A. suum* and 7 dpi for *S.* Typhimurium. The same time points were maintained for coinfected animals and uninfected controls were included at each timepoint.

### Necropsy and Tissue Sampling

Pigs were sedated using ketamine hydrochloride (33 mg/kg, Ursotamin, Serumwerk Bernburg AG, Bernburg, Germany), xylazine (6 mg/kg, Xzlavet, CP-Pharma GmbH, Burgdorf, Germany) and azaperone (4 mg/kg, Stresnil, Janssen-Cilag GmbH, Neuss, Germany) and euthanized by intracardial injection of T61 (10 mg/kg of tetracaine hydrochloride, mebezonium iodide, and embutramide, Intervet Deutschland GmbH, Unterschleißheim, Germany). See Text S1 for tissue sampling protocols.

### *Salmonella* Burden Determination

*Salmonella* burden was determined in cell suspensions obtained from tissue homogenates at different sites (jejunal mLN, ileo-cecal mLN, and spleen). Cell suspensions were plated onto LB plates supplemented with 50 µg/mL nalidixic acid (Carl Roth GmbH) and incubated overnight at 37 °C. The next day *S.* Typhimurium colonies were counted.

### Leukocyte Isolation and Differential Leukocyte Counts

See Text S1.

### Cell Stimulation and Flow Cytometry

For cytokine expression analysis, cells were plated at 1 x 10^6^ cells per well in round bottom 96 well plates and stimulated with 100 ng/mL recombinant porcine IL-12p70 and 100 ng/mL recombinant porcine IL-18 (both from R&D Systems; Minneapolis, MN, USA) for 13 hours at 37 °C and 5% CO_2_ in the presence of Brefeldin A (3 µg/mL; ThermoFisher; Waltham, MA, USA) for the last 10 hours of stimulation. Cells were plated at 3 x 10^6^ per well in a round bottom 96 well plate and allowed to rest overnight at 37°C for T cell cytokine staining. Cells were then stimulated with Phorbol myristate acetate (PMA; 50 ng/mL, Sigma-Aldrich) plus ionomycin (500 ng/mL, Sigma-Aldrich) for 3.5h in presence of Brefeldin A (1 µg/mL, eBioscience) during the last 3h of restimulation.

To assess cellular markers, cells were plated at 2 x 10^6^ cells per well in conical 96 well plates, blocked with mouse serum (1:500), and stained for surface and intracellular markers using the antibodies listed in Table S1 for flow cytometry analysis (BD FACS Canto II, BD FACSARIA III, Flowjo version 10, all BD Life Sciences, Franklin Lakes, NJ, USA) following standard protocols (66). Intranuclear and intracellular marker were stained after fixation and permeabilization of cells with the FoxP3/Transcription Factor staining buffer set (Thermo Fisher) or Cytofix/Cytoperm (BD Biosciences), respectively.

### Histological Analysis

Formalin-fixed tissues were embedded in paraffin, cut into sections using a microtome, and stained with hematoxylin and eosin for histological analysis. See Text S1 for histological scoring.

### Gene Expression Analysis

Frozen tissue samples were processed for RT-qPCR analysis using the InnuPrep RNA Mini Kit (Analytik Jena, Jena, Germany) according to manufacturer’s instructions (See Text S1).

### In Vitro *Salmonella* Infection Assay

See Text S1 for in vitro *Salmonella* infection assay.

### Statistical Analysis

Statistical analysis and visualization of data were performed with GraphPad Prism software (version 9, Dotmatics, Boston, MA, USA) (see Text S1).

## Acknowledgements

We thank Beate Anders, Marion Müller, Christiane Palissa, Bettina Sonnenburg, and Yvonne Weber for their excellent technical support and Dr. Anne Winkler for her design of the graphical abstract and experimental outlines (all from Institute of Immunology, Freie Universität Berlin). We also thank Kristina Dietert, Laura Koschnitzki, Franzisca Möbus, Charline Podehl, Sebastian Scheunemann, and Dominic Schumacher for their essential help with animal care (all from the Veterinary Centre for Resistance Research, Freie Universität Berlin).

## Author Contributions

**Conceptualization:** AM, JSB, & SH. **Formal Analysis:** AM, LO, RMM, ZDM, AAK, RK, SR. **Funding acquisition:** SH. **Investigation:** AM, LO, JSB, AL, RMM, ZDM, PH, AK, RH, JA, SG, MT, LEE, AAK, & RK. **Methodology:** AM, KT, & SH. **Project administration:** AM, JSB, & SH. **Resources:** KT & SH. **Supervision:** JSB, SR, & SH. **Visualization:** AM, ZDM, & SR. **Writing – original draft:** AM. **Writing – review and editing:** AM, LO, JSB, AL, RMM, ZDM, AK, JA, LEE, AAK, KT, SR, & SH.

## Text S1. Supplemental materials and methods

### Tissue Sampling

The trachea was clamped and the lungs were removed. Then, one lung was flushed with phosphate-buffered saline (PBS) supplemented with 2 mM ethylenediaminetetraacetic acid (EDTA) to obtain broncheo-alveolar lavage (BAL) cells. BAL samples were filtered through a 70 µm cell strainer and stored on ice.

Tissue samples for cellular phenotyping from the lung, liver, spleen, jejunum, ileum, and mesenteric lymph nodes (mLN) were collected in wash medium (Roswell Park Memorial Institute 1640 medium with 1% fetal calf serum (FCS), 100 U/mL penicillin, 100 µg/mL streptomycin; all PAN-Biotech GmbH, Aidenbach, Germany) and stored on ice for further processing. Tissue samples for histological analysis were stored in formalin (Roti®-Histofix 10%, Carl Roth GmbH) overnight at room temperature before being transferred to fresh formalin and stored at 4 °C until further processing. Tissue samples for gene expression analysis were snap frozen in liquid nitrogen before being transferred for long term storage at -80 °C.

### Leukocyte Isolation

Peripheral blood mononuclear cells (PBMC) were isolated from blood collected by heart puncture in EDTA coated tubes (S-Monovette®, Sarstedt AG & Co. KG, Nümbrecht, Germany). Blood was diluted 1:2 in 0.9% NaCl and subjected to density gradient centrifugation using Pancoll human solution (density 1.077 g/mL, PAN-Biotech). Splenic and lymph node leukocytes were isolated by mechanical disruption of tissue samples and passage through a 70 µm cell strainer. Lung and liver leukocytes were isolated from multiple tissue pieces which were mechanical homogenized, pooled, and pre-digested using the Lung and Liver Dissociation Kits for mice and the gentleMACS™ Dissociator (Milteny Biotec, Bergisch Gladbach, Germany) using the manufacturer’s protocol with 3X enzyme concentrations.). Small intestinal lamina propria leukocytes were isolated from tissue samples of approximately 10 cm length from the jejunum and ileum after removal of fat, muscle, and connective tissue. Tissues were homogenized and digested with 5 mg/mL Liberase TM, 5 mg/mL Liberase DH, and 4 mg/mL DNaseI (All Roche Diagnostics GmbH, Mannheim, Germany) at 37 °C under gentle agitation (2 x 20 min) prior to mechanical disruption by passage through a 190 µm mesh. All cell suspensions were washed with wash medium and treated with erythrocyte lysis buffer (0.01 M KHCO_3_, 0.155 M NH_4_Cl, 0.1 mM EDTA, pH 7.5) before being resuspended in complete culture medium (cIMDM; Iscove’s Modified Dulbecco’s Medium supplemented with 10% FCS, 100 U/mL penicillin, 100 µg/mL streptomycin, all PAN-Biotech).

### Differential Leukocyte Counts

Leukocyte counts were performed on blood smears and cytospins of BAL fluid cells. Cells on microscope slides were fixed using ethanol and propanol (ROTI®Fix spray, Carl Roth GmbH) then Romanowsky stained (DiffQuick, Labor + Technik, Eberhard Lehmann GmbH, Berlin, Germany). 200 leukocytes were counted and classified to determine percentages of lymphocytes, neutrophils, eosinophils, basophils, and macrophages/monocytes.

### Histological Scoring

For liver tissue scoring was performed according to Con A-induced hepatitis scoring involving the summation of scores for lobular and portal inflammation and necrosis (67). For lung tissue scoring was done according to BCG infection scoring (68). For jejunum and ileum, scoring was carried out according to small intestinal inflammation scoring from Erbet et al. (69). For tissue eosinophil counts, scoring was as follows: 0, normal (physiologic conditions); 1, minimally increased; 2, mildly increased; 3, moderately increased; 4, markedly increased.

### Gene Expression Analysis

Frozen tissue samples were processed for RT-qPCR analysis using the InnuPrep RNA Mini Kit (Analytik Jena, Jena, Germany) according to manufacturer’s instructions. Extracted RNA was transcribed to cDNA using the High Capacity RNA-to-cDNA kit from Applied Biosystem (ThermoFisher; Waltham, MA, USA). Amplification and detection were performed in 96-well optical plates with SYBR-green (Both Applied Biosystems, ThermoFisher). Amplifications were performed in duplicate in a final volume of 20 µL containing 10 µL of 2X SYBR Green I Master Mix and 5 µM of each primer (Table S2). mRNA expression was normalized to the housekeeping gene glycerinaldehyd-3-phosphat-dehydrogenase (*GAPDH*) and standard curves were generated to calculate the efficacy of each primer pair. Relative expression was calculated using the 2^-ΔΔCT^ method (70).

### In Vitro *Salmonella* Infection Assay

BAL cells from uninfected pigs were plated at approximately 3 x 10^5^ cells per well in 24-well cell culture plates and incubated for 1 h at 37 °C. After 1 h, the cells were washed and replaced with fresh cIMDM, leaving the alveolar macrophages attached to the plate which were incubated overnight. The next day, cells were either left untreated, treated with 100 µg/mL IFN-γ, or 10 µg/mL of both IL-4 and IL-13 (IFN-γ and IL-4, R & D Systems; IL-13, Kingfisher Biotech, Saint Paul, MN, USA). Two days later cells were infected with *S.* Typhimurium (ATCC 14028 gyrA (D87Y); strain 8642/KT8640 (pFPV25.1)) grown in aeration at 37 °C at a multiplicity of infection of 10. After addition of bacteria to cell cultures, plates were centrifuged at 150 x *g* for 10 minutes and incubated at 37 °C for 50 mins followed by a change of cell culture medium to cIMDM supplemented with gentamicin 50 µg /mL (PAN-Biotech) and further incubation for 1 h. Medium was then changed to cIMDM with gentamicin 10 µg /mL and incubated overnight (total infection time ∼ 16 h). The next day, cells were analyzed by microscopy and imaged on an Axiovert.A1 microscope equipped with a Colibri 7 solid-state light source and Axiocam 503 mono microscope camera using Zen (v 2.3) software (Carl Zeiss, Oberkochen, Germany). Cells were then washed with PBS and removed from the cell culture plates using trypsin-EDTA (PAN-Biotech) prior to staining with fixed viability dye and analysis by flow cytometry.

### Statistical Analysis

Statistical analysis and visualization of data were performed with GraphPad Prism software (version 9, Dotmatics, Boston, MA, USA). Data were tested for normality using the Shaprio-Wilk test. Parametric data were analyzed by one-way analysis of variance followed by Tukey’s multiple comparisons test. Non-parametric data were analyzed by the Kruskal-Wallis test followed by Dunn’s multiple comparisons test. Differences were considered statistically significant at *p* < 0.05 and indicated as follows: *p* ≤ 0.05 (*), *p* ≤ 0.01 (**), *p* ≤ 0.01 (***), and *p* ≤ 0.001 (****).

---

**Table S1.**
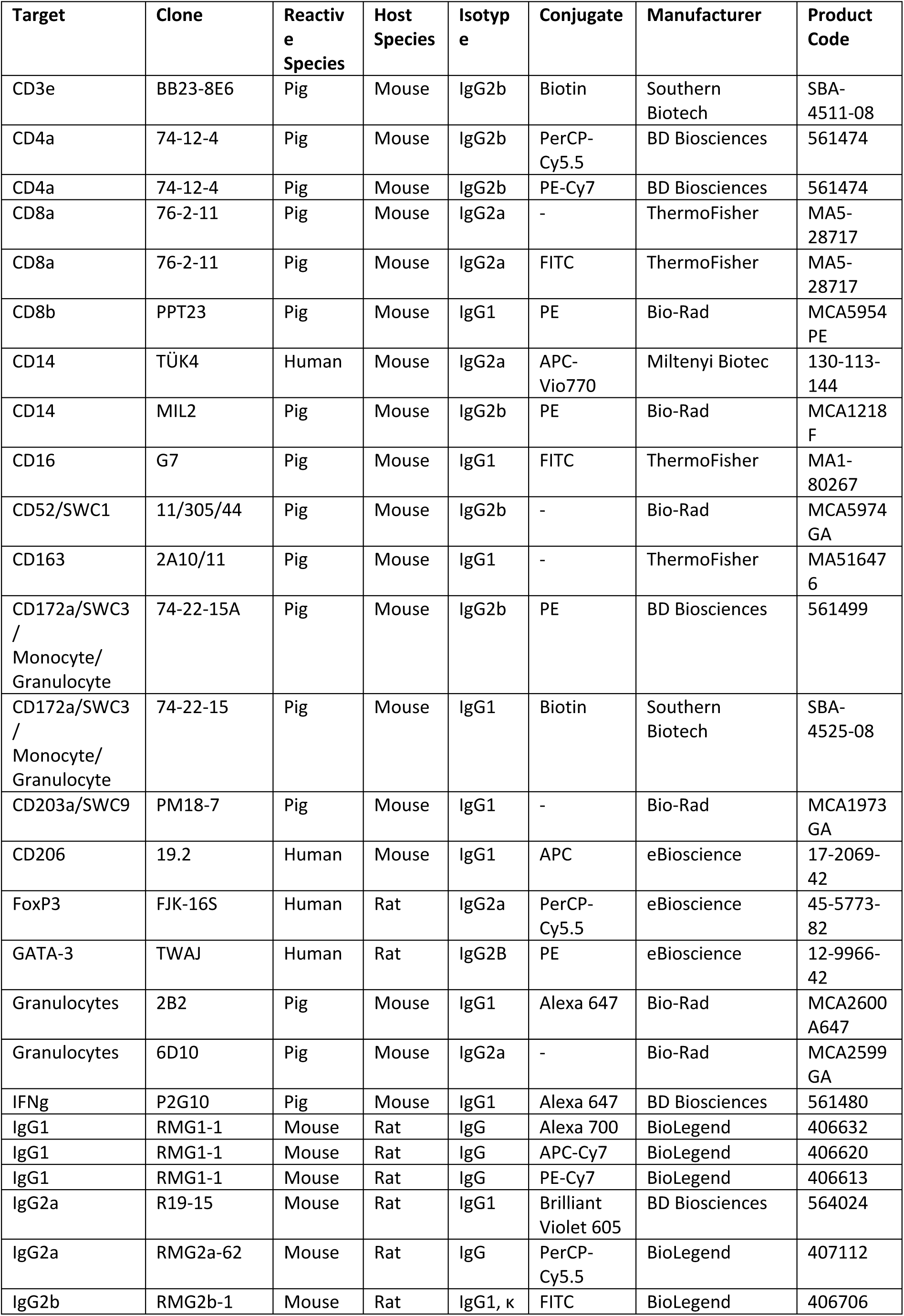
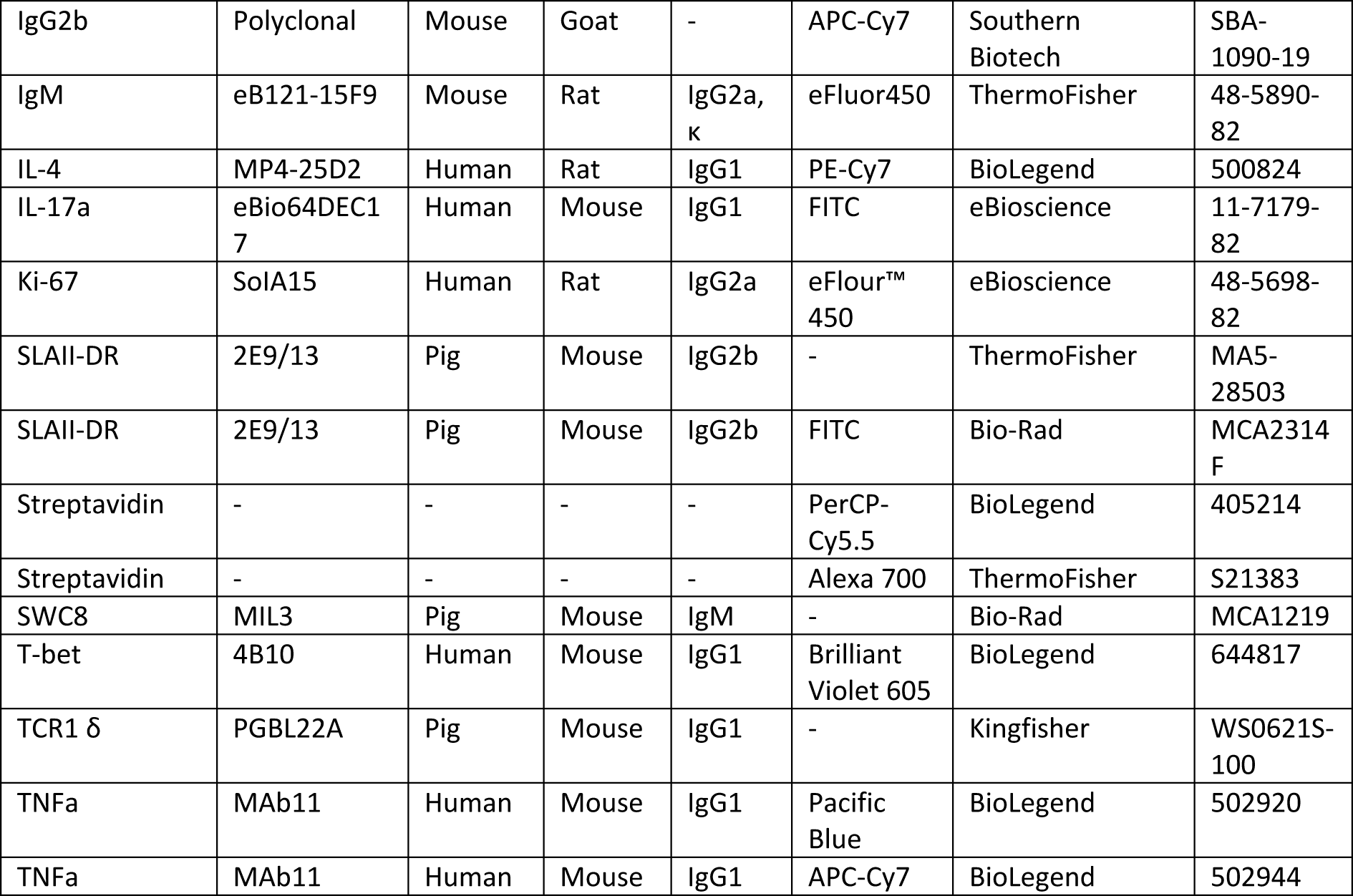
Antibodies used in this study

**Table S2.**
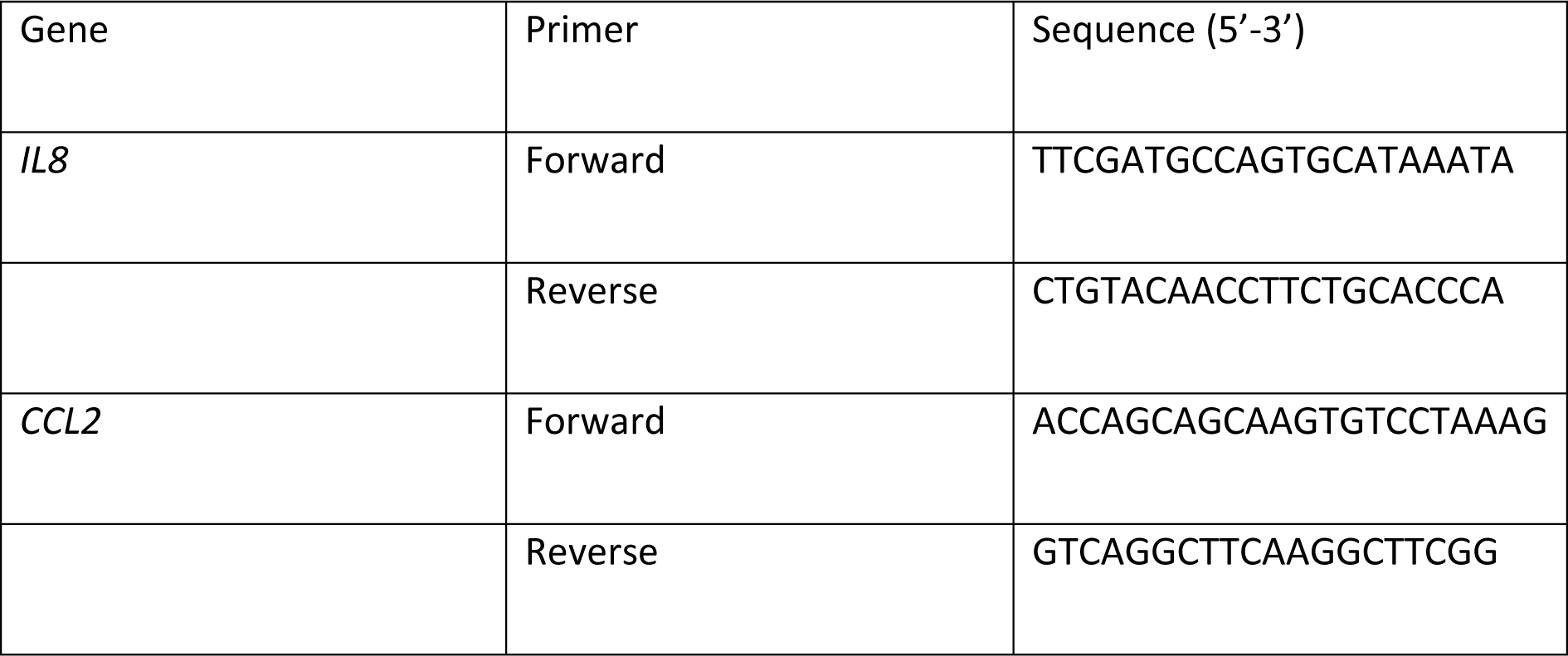
Primers used in this study

**Figure S1.**
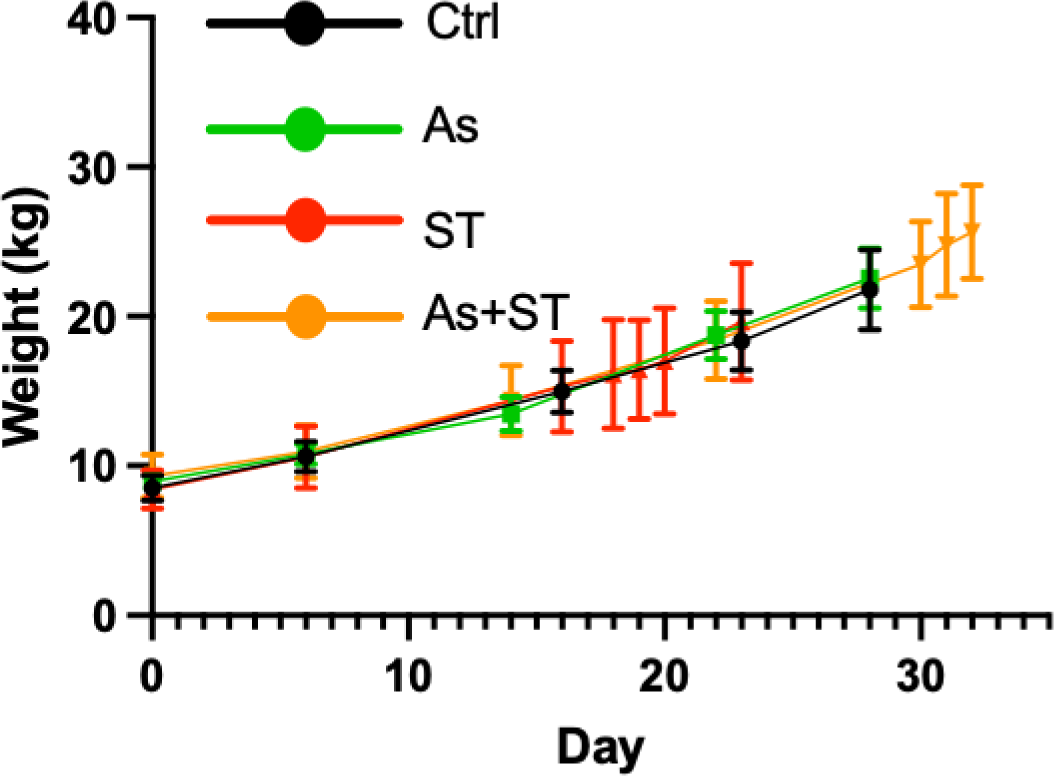
Infections had no impact on pig weight gain. Shown are average weights (kg) ± SD across groups throughout the experiment.

**Figure S2.**
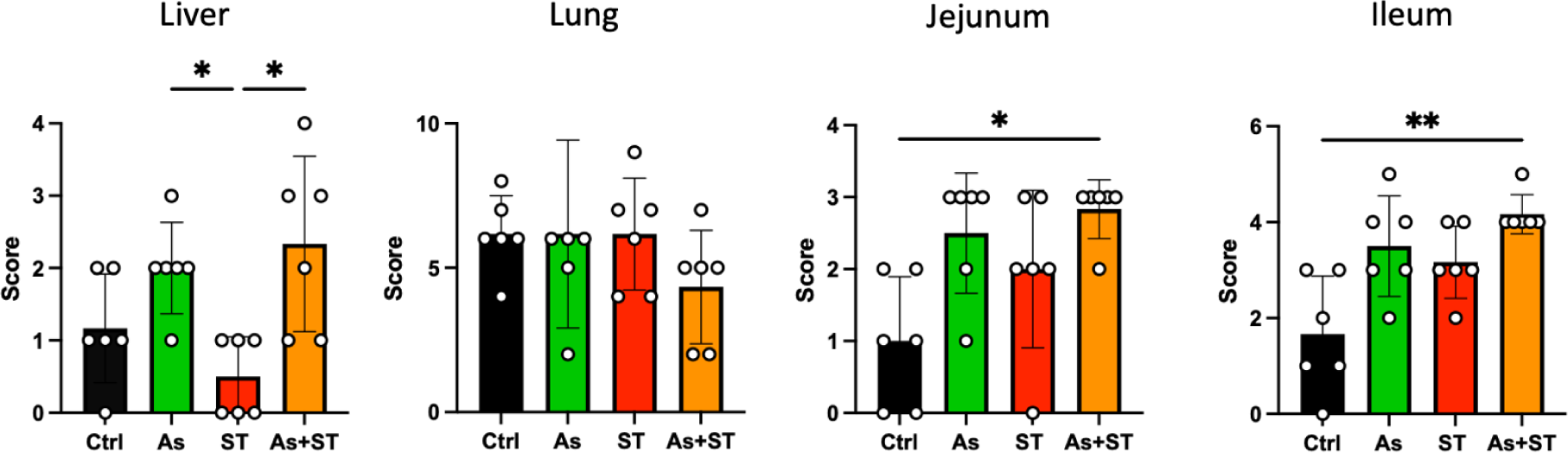
Histopathology scoring of tissues impacted by *Ascaris* and *Salmonella*. Pathology scores determined as described in the methods section. Statistical significance was determined by one-way analysis of variance followed by Tukey’s multiple comparisons test, * *p* ≤ 0.05, ** *p* ≤ 0.01.

